# Integration of spatial transcriptomic and single cell sequencing identifies expression patterns underlying immune and epithelial cell cross-talk in acute kidney injury

**DOI:** 10.1101/2021.01.19.427258

**Authors:** Ricardo Melo Ferreira, Angela R. Sabo, Seth Winfree, Kimberly S. Collins, Danielle Janosevic, Connor Gulbronson, Ying-Hua Cheng, Lauren Casbon, Daria Barwinska, Michael J. Ferkowicz, Xiaoling Xuei, Chi Zhang, Kenneth W. Dunn, Katherine J. Kelly, Timothy A. Sutton, Takashi Hato, Pierre C. Dagher, Tarek M. El-Achkar, Michael T. Eadon

**Author notes:** Correspondence information: Michael Eadon, MD, 950 West Walnut Street R2, 202, Indianapolis, IN 46202, Tarek El-Achkar, MD, 950 West Walnut Street R2, 202, Indianapolis, IN 46202.

## Abstract

Despite important advances in studying experimental and clinical acute kidney injury (AKI), the pathogenesis of this disease remains incompletely understood. Single cell sequencing studies have closed this knowledge gap by characterizing the transcriptomic signature of different cell types within the kidney. However, the spatial distribution of injury can be regional and affect cells heterogeneously. We first optimized coordination of spatial transcriptomics and single nuclear sequencing datasets, mapping 30 dominant cell types to a human nephrectomy sample. The predicted cell type spots corresponded with the underlying hematoxylin and eosin histopathology. To study the implications of acute kidney injury on the distribution of transcript expression, we then characterized the spatial transcriptomic signature of two murine AKI models: ischemia reperfusion injury (IRI) and cecal ligation puncture (CLP). Localized regions of reduced overall expression were found associated with tissue injury pathways. Using single cell sequencing, we deconvoluted the signature of each spatial transcriptomic spot, identifying patterns of colocalization between immune and epithelial cells. As expected, neutrophils infiltrated the renal medullary outer stripe in the ischemia model. Atf3 was identified as a chemotactic factor in S3 proximal tubule cells. In the CLP model, infiltrating macrophages dominated the outer cortical signature and Mdk was identified as a corresponding chemotactic factor. The regional distribution of these immune cells was validated with multiplexed CO-Detection by inDEXing (CODEX) immunofluorescence. Spatial transcriptomic sequencing can aid in uncovering the mechanisms driving immune cell infiltration and allow detection of relevant subpopulations in single cell sequencing. The complementarity of these technologies facilitates the development of a transcriptomic kidney atlas in health and disease.

## Introduction

Acute kidney injury (AKI) is a devastating disease with negative impact on morbidity and mortality. Developing therapeutic targets to treat AKI requires a better grasp of its molecular pathogenesis. Despite important advances in understanding this disease, the pathogenesis of AKI at the cellular and molecular levels remains incompletely understood. This is partially due to the diverse renal milieu of heterogeneous cell types (epithelial, endothelial, fibroblast, vascular smooth muscle, resident immune, and infiltrating immune cells) that interact with each other within a cosmos of unique microenvironments. Furthermore, AKI differentially affects the kidney’s diverse array of cells, which increases the complexity of disease (1). Recently, single cell and single nuclear sequencing have proved major breakthroughs in the creation of a molecular atlas of the kidney (2–5) by defining the transcriptomic signatures of specific cells within the kidney. However, spatial anchoring is essential to defining the relationship between cells and structures within specific renal microenvironments.

Single cell and single nuclear sequencing afford indirect spatial localization. In contrast, spatial transcriptomics platforms enable measurement of whole transcriptome mRNA expression of thousands of genes superimposed upon histological information from the same tissue section. Gene expression profiles are mapped back to their original location, enabling a direct link between histology and gene expression (6). Integration of the single cell sequencing with spatial transcriptomics (ST) improves power and enables *in situ* visualization of signatures with mapping of a greater number of cell types than spatial transcriptomics alone (7). Together, the two orthogonal datasets allow determination of where immune cells reside in proximity to other cells in disease states and how regional injury influences the signature of epithelial cells.

In the present study, we utilized single nuclear and single cell sequencing datasets to map cell types back to spatial transcriptomic anchoring landmarks (spots) overlaid upon human and murine kidney. We first optimized the methodology in the human kidney. We then examined differentially expressed genes (DEGs) and pathways in murine ischemia-reperfusion injury (IRI) and cecal ligation puncture (CLP) models of AKI. We used the transcriptomic signatures derived from the Visium 10x Spatial Gene Expression platform to co-localize immune cells with epithelial cells, defining the distribution of immune cell transcript expression in these models. Key chemotactic factors expressed in epithelial cells were identified which contribute to immune cell infiltration and cross-talk in each injury model. Immune cell type distributions were validated with spatially resolved multiplexed CO-Detection by inDEXing (CODEX) immunofluorescence. Together, the complementary single cell and spatial transcriptomic datasets are synergistic, enabling the development of a transcriptomic atlas of the kidney in health and disease, with direct correlation to histopathology. These tools will enhance the diagnostic capability of renal pathologists interpreting manifestations of AKI in kidney tissue.

## Results

### Unsupervised mapping and cell type identification within the spatial transcriptomic map of the human kidney

We sought to map transcriptomic signatures directly upon a histologic section from a human reference nephrectomy stained with hematoxylin and eosin (H+E). The tissue was obtained from a 59 year-old female with minimal glomerular obsolescence and interstitial fibrosis (each affecting less than 10% of the glomeruli or renal parenchyma respectively). No arteriolar hyalinosis was observed. An Optimal Cutting Temperature (OCT) compound embedded tissue section underwent H+E staining and microscopy, followed by permeabilization, RNA isolation, and sequencing with spatially localized barcodes to map back the transcriptomic signature directly upon the histologic image.

The result of the spatial transcriptomic (ST) mapping is a set of “spots” (55 μm in diameter), each with its own expression signature. These spots are overlaid upon the histologic image of the kidney based on the localization barcodes (Figure 1A-B) and are clustered in an unsupervised fashion according to the gene expression of each spot. The identity of these clusters was established and named according to known kidney regions and cell types using differentially expressed marker genes and the underlying histology. Nine unsupervised clusters were generated by Space Ranger (Figure 1B), with expression signatures aligning with known marker genes of glomeruli and various tubular subsegments. These clusters were overlaid upon histological features including glomeruli, tubules, vascular structures, and medullary rays, which are readily observed. On average, 10,270 counts were measured and 3,205 genes were detected per spot. The average number of unique genes detected was 17,506 per cluster, with at least 16,000 detected in every cluster.

**Figure 1:**
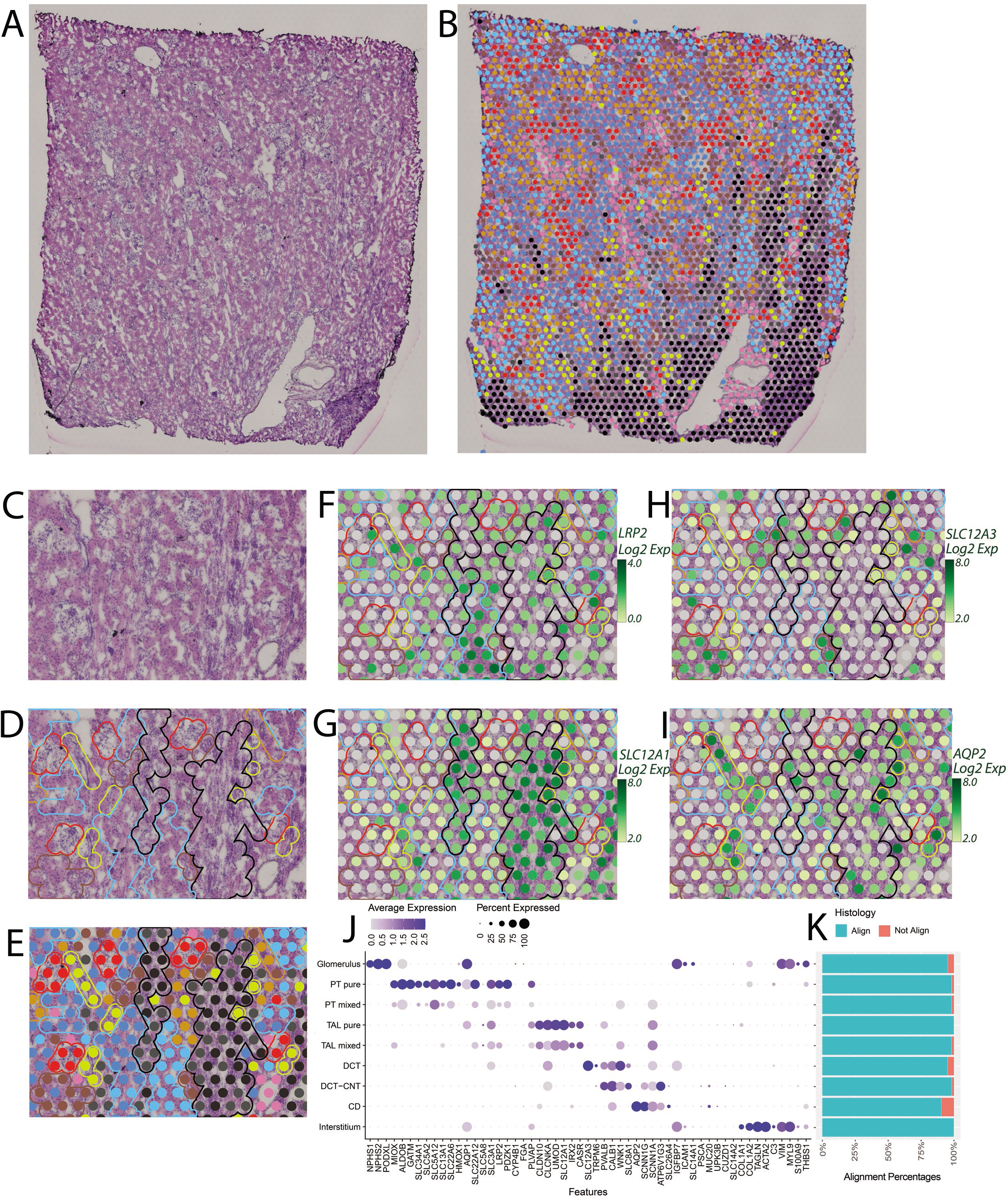
Spatial Transcriptomics of Human Biopsy. (**A**) Hematoxylin and Eosin staining of the human reference nephrectomy. (**B**) The nine Spatial Transcriptomics unsupervised clusters are overlaid upon the nephrectomy. Glomeruli can be seen scattered across the cortex in red. A medullary ray is seen in the right lower quadrant of the sample. Midsized vessels are often overlaid by pink interstitial cluster spots. Pure clusters are defined as located mainly over the associated structure, while Mixed clusters present some overlap with neighboring structures. (**C**) Close zoom showing histological structures of reference nephrectomy. (**D**) Highlight of histological structures in nephrectomy. (**E**) Unsupervised clusters over nephrectomy zoom with highlights. (**F-I**) Expression level of LRP2, SLC12A1, AQP2 and SLC12A3 in the spots over the zoom section with histological feature highlighted. (**J**) Dotplot presenting expression of markers used to classify unsupervised clusters. (**K**) In 5 random fields covering 40.0% of all spots, the histology underlying each spot was assessed and the percentage of concordance is provided. All clusters held greater than 90% concordance with their corresponding histology. (Abbreviations: PT – Proximal Tubule; S1, S2, S3 – Segments of proximal tubule; TAL – Thick Ascending Limb; DCT – Distal Convoluted Tubule; CNT – Connecting Tubule; CD – Collecting Duct). Each spot is 55 μm in diameter.

Spots assigned to a given cluster consistently localized over the expected underlying histology. An example of cluster mapping and identification is provided in Figure 1C-I. A subset of marker genes is provided in Figure 1J with a full set given in Supplemental Table S1. A close-up of the H+E section is provided without (Figure 1C) and with (Figure 1D-E) unsupervised clusters overlaid. Glomeruli, small vessels, tubules, and a portion of the medullary rays can be visualized. Transcriptomic spots associated with glomerular genes (red) clearly overlie histologically identified glomeruli. The interstitial cluster which includes expression of fibroblast, vascular smooth muscle cell (VSMC), and endothelial cell marker genes is seen located over vessels (pink). Although tubular morphology in OCT embedded sections is imperfect, distinctions can still be observed. The spatial transcriptomic defined proximal tubule clusters overlie brightly eosinophilic tubules without visible lumens. This histologic pattern corresponded with megalin (*LRP2*) expression (Figure 1F). Spots assigned to thick ascending limb (TAL) clusters are congregated over medullary rays and correlate with expression of the loop diuretic sensitive sodium-potassium 2-chloride transporter (*SLC12A1*, Figure 1G). Distal convoluted tubules (DCTs) and connecting tubules are characterized by eosinophilic tubules with open lumens. DCTs had high corresponding expression of the thiazide sensitive sodium-chloride transporter (*SLC12A3*, Figure 1H). Collecting ducts can be identified histologically with their characteristic nuclei in cuboidal cells and reduced eosinophilia. These tubules were overlaid by spots with high aquaporin-2 (*AQP2*) expression (Figure 1I).

Since each ST spot is 55 μm in diameter, roughly the size of a tubular cross section, the spots can overlie multiple cell types. Two PT and two TAL clusters were obtained in the unsupervised mapping. One PT and one TAL cluster each had stronger enrichment of known marker genes and were labeled “pure”. The clusters with less enrichment of marker genes or signatures with elements of two or more cell types were labeled “mixed”. The mixed TAL signature was seen on the periphery of the medullary rays, and the pure cluster was more centrally located (Figure 1B).

As stated, spots were classified into clusters based on the known gene expression markers of renal cell types (Figure 1J, Supplemental Table S1). The spots classified based on their gene expression signature mapped to expected histological/renal structures consistently. We assessed the consistence of the assigned transcriptomic cluster to the underlying histology in five randomly selected fields (Figure 1K). The overall correlation between assigned clusters and histologic structures was 97.6%. All clusters mapped to a corresponding histologic structure at greater than 95% accuracy except the collecting duct (90.7%) which had the fewest overall spots in the specimen. The most frequent reason for discordance was the lack of significant tissue underlying a spot (e.g. a spot at the edge or within a large vessel).

### Mapping of single nuclear sequencing clusters to human kidney tissue

A UMAP of the unsupervised clusters obtained through Space Ranger is provided in Figure 2A. To improve the specificity of spot mapping and identify less frequent cell types which may contribute to the signature of a spot, we re-clustered a publicly available single nuclear RNA sequencing (snRNAseq) dataset from human kidney samples (Figure 2B) (2). The full list of DEG markers of each scRNAseq cluster are found in Supplemental Table S2. Thirty cell type clusters were identified. In a process similar to single cell cluster integration (8), transfer scores were assigned to ST spots for each of the 30 snRNAseq clusters based on anchoring of common neighbors. Each spot is then labelled according to the snRNAseq cluster with the highest transfer score. We then deconvoluted the unsupervised ST clusters according to the transferred labels from the snRNAseq clusters (Figure 2C). The percentage of spots from the unsupervised ST clusters relabeled according to the snRNAseq clusters was assessed. A strong correlation was found between the unsupervised ST classified spots and the spots redefined with the expected snRNAseq transfer labels.

**Figure 2:**
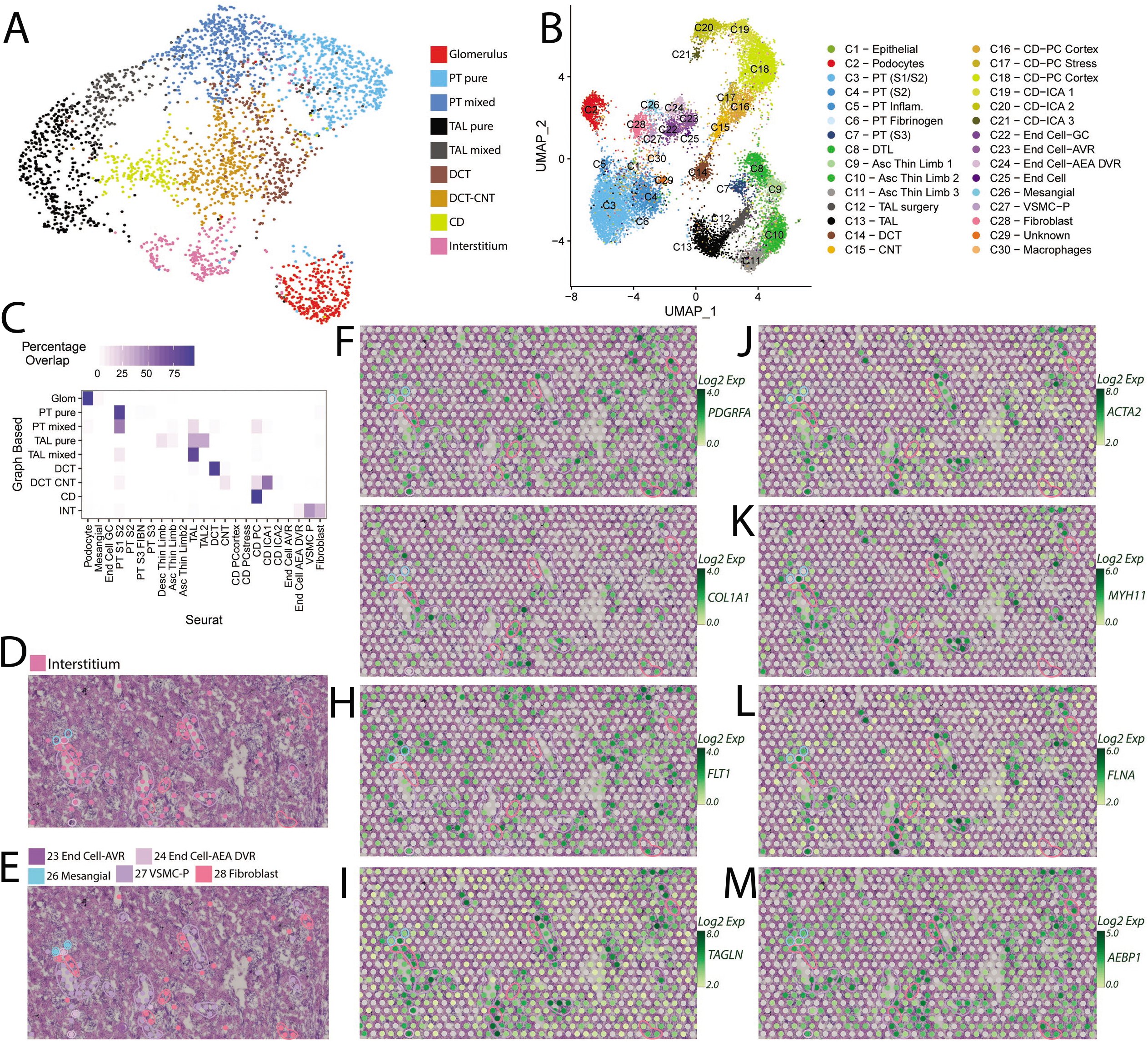
Transfer of Single Nucleus clusters to Spatial Transcriptomics. (A) UMAP projection of Spatial Transcriptomic data with nine unsupervised clusters defined by Space Ranger. Pure cluster spots were frequently located over their corresponding histology, while mixed clusters frequently overlap neighboring structures. (B) UMAP projection of the Single Nucleus data (GSE121862) depicting 30 kidney cell clusters obtained from Pagoda. (C) Heatmap presenting the percentage of unsupervised cluster spots overlapping with single nucleus mapped clusters. Strong correlation is seen between expected clusters. (D) Close zoom of Hematoxylin and Eosin stained human reference nephrectomy with unsupervised interstitium cluster spots and histological structures highlighted. (E) Close zoom of Hematoxylin and Eosin staining of human reference nephrectomy with mapped single nucleus clusters associated with interstitium, and histological structures highlighted. (F-M) Feature plots depict the expression levels of PDGFRA, COL1A1, FLT1, TAGLN, ACTA2, MYH11, FLNA and AEBP1 in the zoomed region with histological features highlighted. (Abbreviations: PT – Proximal Tubule; S1, S2, S3 – Segments of proximal tubule; TAL – Thick Ascending Limb; DCT – Distal Convoluted Tubule; CNT – Connecting Tubule; CD – Collecting Duct; DTL – Descending Thin Limb; Asc – Ascending; PC – Principal Cells; IC – Intercalated Cells; End – Endothelial; GC – Glomerular Capsule; AVR – Ascending Vasa Recta; AEA Afferent and Efferent Arterioles; DVR – Descending Vasa Recta; VSMC-P – Vascular Smooth Cells and Pericytes). Each spot is 55 μm in diameter

Several cell types that were not represented in the unsupervised ST clusters had spots assigned to them, although, since each spot covers multiple cells, the cell type is assigned to the most dominant cell type in that 55 μm spot. For example, multiple cell types contribute to the expression signature of the interstitium (9). The inclusion of snRNASeq data resolved the spots localizing in the Interstitium to a more specific set of cell types (Figure 2D-M). This dataset allowed discrimination of spots assigned to fibroblasts, VSMCs, and some endothelium subtypes. Feature plots of known markers for the Interstitium are shown in Figures 2F to 2M.

### Spatial transcriptomic cell type localization in the murine kidney

Acute kidney injury (AKI) does not affect all regions of the kidney uniformly. The etiology of injury may differentially impact the spatial distribution of gene expression, especially in the early course of disease. Therefore, we applied special transcriptomics on two common murine models of AKI (IRI and CLP), sacrificed at 6 hours. The unsupervised ST cluster mapping is shown in Figure 3 for the sham, IRI and CLP models. In matched cross-sections of the three models, minimal histopathologic injury was observed with H+E staining at the 6 h time point (Figure 3A). At low power magnification, the IRI tissue area is larger (25.5 mm^2^) than the sham (16.9 mm^2^) and CLP (15.9 mm^2^) sections. Figure 3B overlays each section with the clusters obtained from Space Ranger. Two clusters were only present in the IRI model: the interstitium and urothelium clusters, the latter possibly due to the size of the papilla in the section. The three models were merged and normalized to allow comparisons between models (Figure 3C). A full set of DEGs between unsupervised ST clusters is found in Supplemental Table S3. Murine cells and tubules are smaller than their human counterparts, so each 55 μm spot is labeled by its dominant contributor. Fewer unsupervised clusters were identified in the mouse than the human. For example, the DCT, CNT, and CD of the distal nephron all clustered together. Similarly, the PT S3 segment cluster localizes to the outer stripe and contains gene expression from the neighboring TAL. A subset of marker genes used to identify the clusters are depicted in Figure 3D.

**Figure 3:**
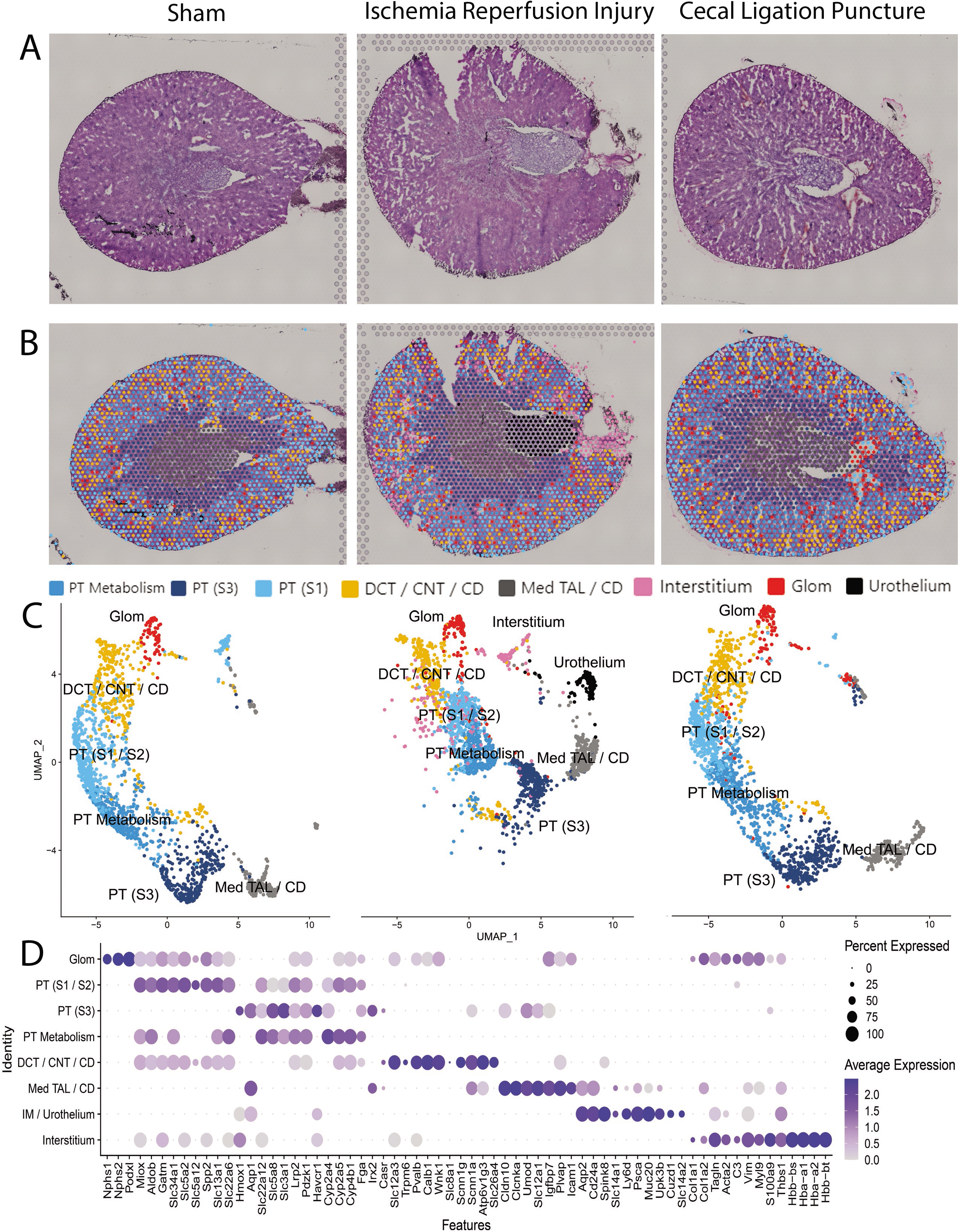
Spatial Transcriptomics of Murine Kidneys. (**A**) Hematoxylin and Eosin staining of sections of three murine models, Sham, Ischemia Reperfusion Injury (IRI) and Cecal Ligation Puncture (CLP), respectively. (**B**) Spatial Transcriptomic cluster spots are overlaid upon each murine kidney. (**C**) UMAP of the spatial clusters after the data was merged, split by tissue of origin. (**D**) Dotplot presenting expression of markers used to classify the spatial clusters. (Abbreviations: PT – Proximal Tubule; S1, S2, S3 – Segments of proximal tubule; Med – Medullary; TAL – Thick Ascending limb; DCT – Distal Convoluted Tubule; CNT – Connecting Tubule; CD – Collecting Duct; Glom - Glomerulus). Each spot is 55 μm in diameter.

### Regional expression differences and pathway enrichment in injury models

To investigate regional expression differences between disease models, we first calculated the overall expression level of each spot, as measured in total read counts (Figure 4A). We then identified regions of the IRI and CLP models that had areas of reduced expression when compared to Sham. These regions of “low” expression in IRI and CLP were selected and compared to size matched regions with relatively “preserved” expression. The low expression regions were also compared to identically sized regions in the sham.

**Figure 4:**
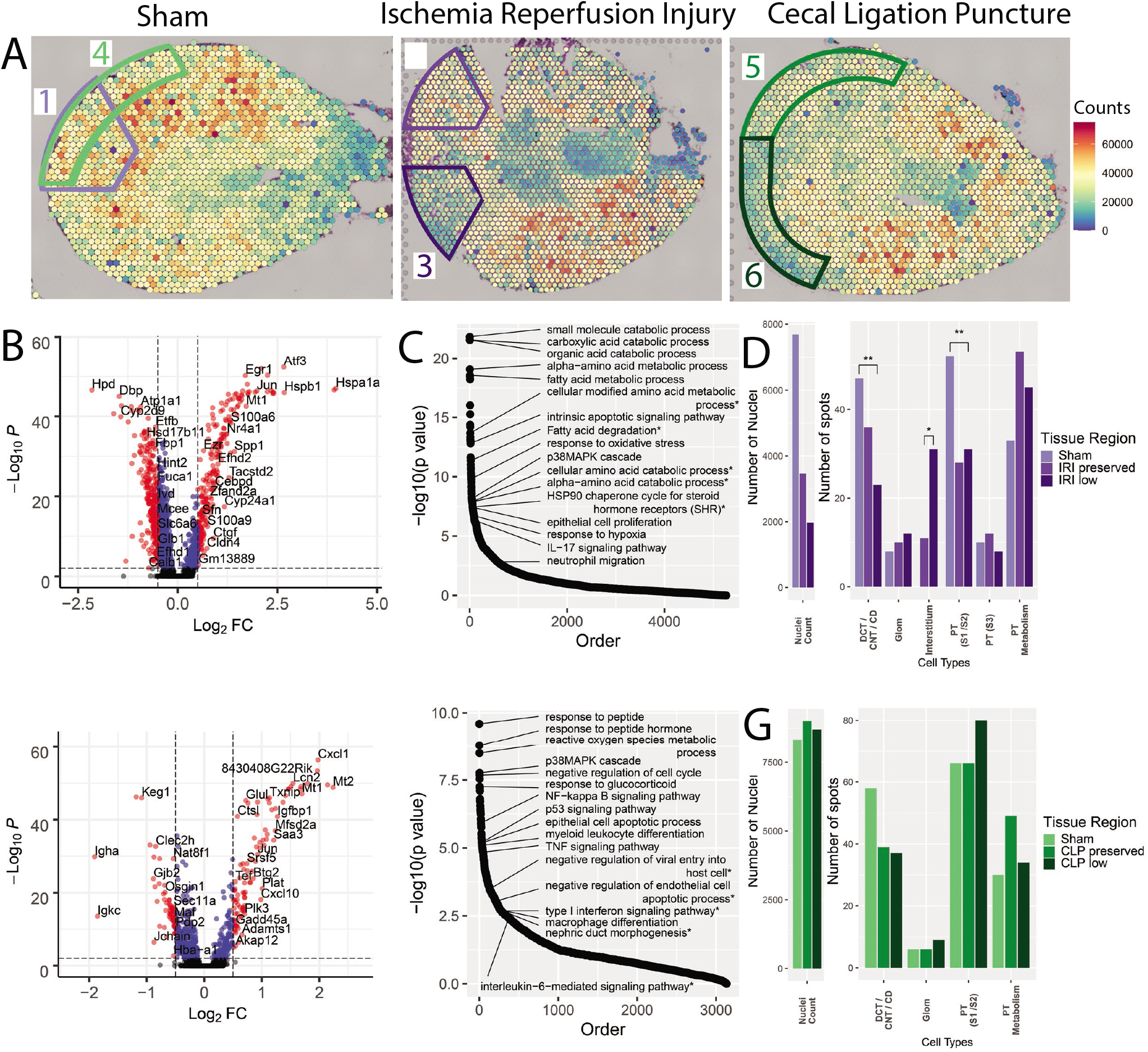
Spatial distribution of Injury on Murine Models. (A) Total expression in read counts was summed for each spatial transcriptomic spot and the total expression level was overlaid upon each of the three murine models: Sham, Ischemia Reperfusion Injury (IRI), and Cecal Ligation Puncture (CLP). Regions of interest and comparator regions are highlighted. In the Sham, areas are selected to serve as reference to IRI (1, outlined in purple) and to CLP (4, outlined in green). In the IRI section, region 2 corresponds to the relatively “preserved” overall expression region and region 3 corresponds to a region of low relative expression. In the CLP section, analogous regions of preserved expression (5) and “low” expression (6) were selected. The regions were defined with similar areas within each comparison. (B) Volcano plot comparing the low expression region in the IRI to the equivalent region in the Sham. Despite the overall reduced expression of the IRI region, many individual genes were up-regulated in IRI (right). (C) Pathways enriched for the Differentially Expressed Genes (DEGs) between the low expression region in IRI when compared to Sham. (D) Bar plots showing the number of nuclei and number of spots of each cluster in the three purple comparison regions. The asterisks indicate the significance level (* - p < 0.1, ** - p < 0.001). (E) Volcano plot comparing the low expression region in the CLP to the equivalent region in the Sham with up-regulated genes in CLP on the right. (F) Pathways enriched for the DEGs between the low expression region in CLP when compared to Sham. (G) Bar plots showing number of nuclei and number of spots of each cluster in the three green comparison regions. Each spot is 55 μm in diameter.

The DEGs, across all spots regardless of cluster identity, were uncovered in the IRI low expression region and the equivalent sham region (Figure 4B). Despite the overall reduction in total read counts in IRI, a number of genes were significantly up regulated in the low expression region. The enriched pathways based on the DEGs are presented in Figure 4C with a subset of pathways annotated. Several enriched pathways were identified related to metabolism of amino acids and fatty acids, possibly related to a metabolic shift associated with hypoxia (10–14). Others pathways suggest prominent injury response mechanisms, such as apoptosis, oxidative stress, and the p38MAPK cascade (15–17). Interleukin-17 signaling and enrichment of neutrophil migration indicate a potential inflammatory response of neutrophils in the IRI model (18–20). The full pathway list is found in Supplemental Table S4 and DEGs are found in Supplemental Table S5.

In a separate comparison, the IRI “low” and “preserved” expression regions were compared (Supplemental Table S4 and S5). Similar DEGs and pathways were identified, albeit at lower levels of significance. The IRI “preserved” expression region fell along the spectrum of expression changes between the IRI “low” expression and the sham regions. We assessed the total nuclei count and spot distribution in each of these selected regions (Figure 4D). The sham region had approximately twice the nuclei of the IRI “normal” region and four-fold more nuclei than the IRI “low” region. This finding is consistent with the overall changes in total regional expression. We speculate that this relative loss of nuclei in the IRI regions is due to a combination of interstitial edema and apoptosis, which is not surprising at the early stages of this type of injury (6 hours after IRI). Furthermore, the change in the distribution of spot identity is also supportive, wherein the IRI model had an increased number of spots assigned to the interstitial cluster compared to the sham region, which had no spots mapping to the interstitial cluster. A reduced number of epithelial cells in the spots of the injured tissue may account for this shift in distribution. Indeed, fewer spots were found to map to tubular clusters (DCT / CNT / CD and PT (S1 / S2)) in the IRI model.

An equivalent analysis was performed for the CLP model (Supplemental Table S4 and S5). The DEGs and enriched pathways between the CLP “low” and sham regions were consistent with known changes in the CLP model, including p53 signaling, cell cycle arrest, apoptosis, TNF signaling, and macrophage differentiation (Figure 4E-F). The “preserved” CLP region had muted expression differences and pathway enrichment, again falling along the differential expression spectrum between the CLP “low” and sham comparison. In contrast to the IRI model, differences in the nuclei count and the spot cluster identity distribution were not found (Figure 4G). Together, the expression patterns, nuclei counts, and similarity in spot identity distribution between regions in the CLP model suggest a very different injury pattern than IRI. As compared to IRI, CLP injury may lead to less tubular cell apoptosis or space occupying interstitial edema.

### Mapping of single cell sequencing clusters to murine kidney tissue

To improve resolution and cell type specificity, murine single cell RNA sequencing (scRNAseq) datasets were clustered to build a common reference murine kidney cell atlas across models. To define cell types relevant to each model, kidneys of animals from the same sham and IRI models were disaggregated for scRNAseq. To obtain signatures of cell types relevant to the CLP model, publicly available scRNAseq data from an endotoxin lipopolysaccharide (LPS) model (sacrificed at 4 hours post endotoxin) and a corresponding sham animal were utilized (21). Although the CLP and LPS models are not identical, the scRNAseq dataset serves a broader purpose by allowing the identification of cell type signatures within the spatial transcriptomic samples. Expected epithelial, infiltrating immune, and other cell types were identified based on their expression signature (Figure 5A, Supplemental Table S6). The common scRNAseq clusters from the composite UMAP of all 4 animals (IRI and sham, LPS and sham) were then mapped upon the histologic section of each model (Figure 5B-E). Beyond the traditional epithelial cell types, we identified two clusters of injured proximal tubule cells highly expressing either Havcr1 (Kim1) or Fibrinogen (22). The cluster expressing Havcr1, PT (S3-OS), also expresses markers of the S3 proximal tubule, suggesting injury to the outer stripe region of the medulla. The PT Fibrinogen cluster also had some expression of Havcr1, but was rich in Fga, Fgb and Fgg and markers of the more cortical S1 and S2 PT markers.

**Figure 5:**
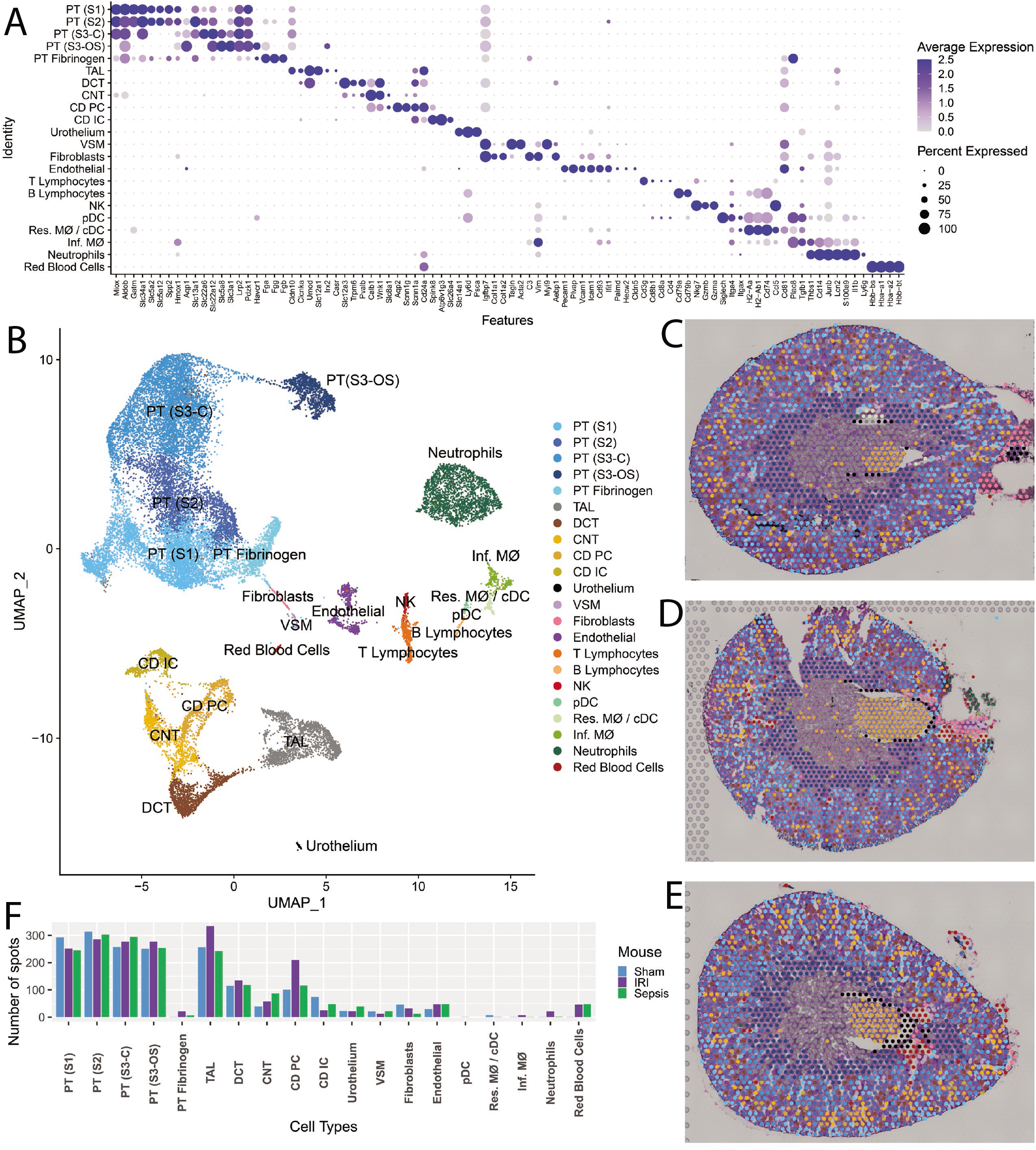
Single Cell murine data and cluster transfer to Spatial Transcriptomics. (**A**) Dot plot presenting expression levels of markers used to define clusters in the single cell data. This dataset consists of four murine samples, an Ischemia Reperfusion Injury (IRI) mouse with corresponding sham and a Lipopolysaccharide endotoxin administered mouse with its corresponding sham. (**B**) UMAP presenting the clusters obtained from the single cell data. (**C-E**) Mapping of the single cell clusters over the three murine spatial transcriptomics sections: Sham, IRI and Cecal Legation Puncture, respectively. (**F**) Bar plot presenting the number of spots mapped to each of the single cell clusters. (Abbreviations: PT – Proximal Tubule; S1, S2, S3 – Segments of proximal tubule; S3-C – Cortical section of S3; S3-OS – Outer Stripe section of S3; TAL – Thick Ascending limb; DCT – Distal Convoluted Tubule; CNT – Connecting Tubule; CD – Collecting Duct; PC – Principal Cells; IC – Intercalated Cells; VSM – Vascular Smooth Muscle; NK – Natural Killer Cells; pDC - Plasmacytoid Dendritic Cells; cDC - Conventional Dendritic Cells; Res. MΦ – Resident Macrophages; Inf. MΦ – Infiltrating Macrophages). Each spot is 55 μm in diameter.

We examined differential expression between the IRI and CLP models in a subset of epithelial cell clusters in the ST dataset (Supplemental Figure S1) and between the IRI and LPS scRNAseq datasets (Supplemental Figure S2). A high degree of overlap was found between the DEGs identified in each technology’s comparison for epithelial cells, but not for immune cells due to the paucity of ST immune cell spots.

Among the immune cell clusters mapped using the scRNAseq dataset, we identified neutrophils, natural killer cells, B and T lymphocytes, plasmacytoid dendritic cells (high in Siglech), and two infiltrating macrophage clusters, Ear2+ (Ly6c^Lo^) and Chil3+ (Ly6c^Hi^) (23). A single cluster was found to contain both resident macrophages and classical dendritic cells (high in H2-Aa, H2-Ab1), which could not be sub-clustered due to cell sample size. The resultant composite cluster set was used to transfer the scRNAseq cluster labels to the ST spots.

The number of spots assigned to each cell type is provided in Figure 5F. Although the single cell dataset yielded an abundant immune cell population, very few ST spots were assigned to those cell types. This apparent discord may be related to the dominant transcriptomic signature originating from the more abundant epithelial cells. A transfer score system was developed to understand the rank and relative contribution of each scRNAseq cell type to the signature of a given ST spot (Supplemental Table S7). After suppression of the epithelial and endothelial scRNAseq clusters, the immune cell and fibroblast scRNAseq cluster labels were remapped over the histologic images (Figure 6 and Figure 7).

**Figure 6:**
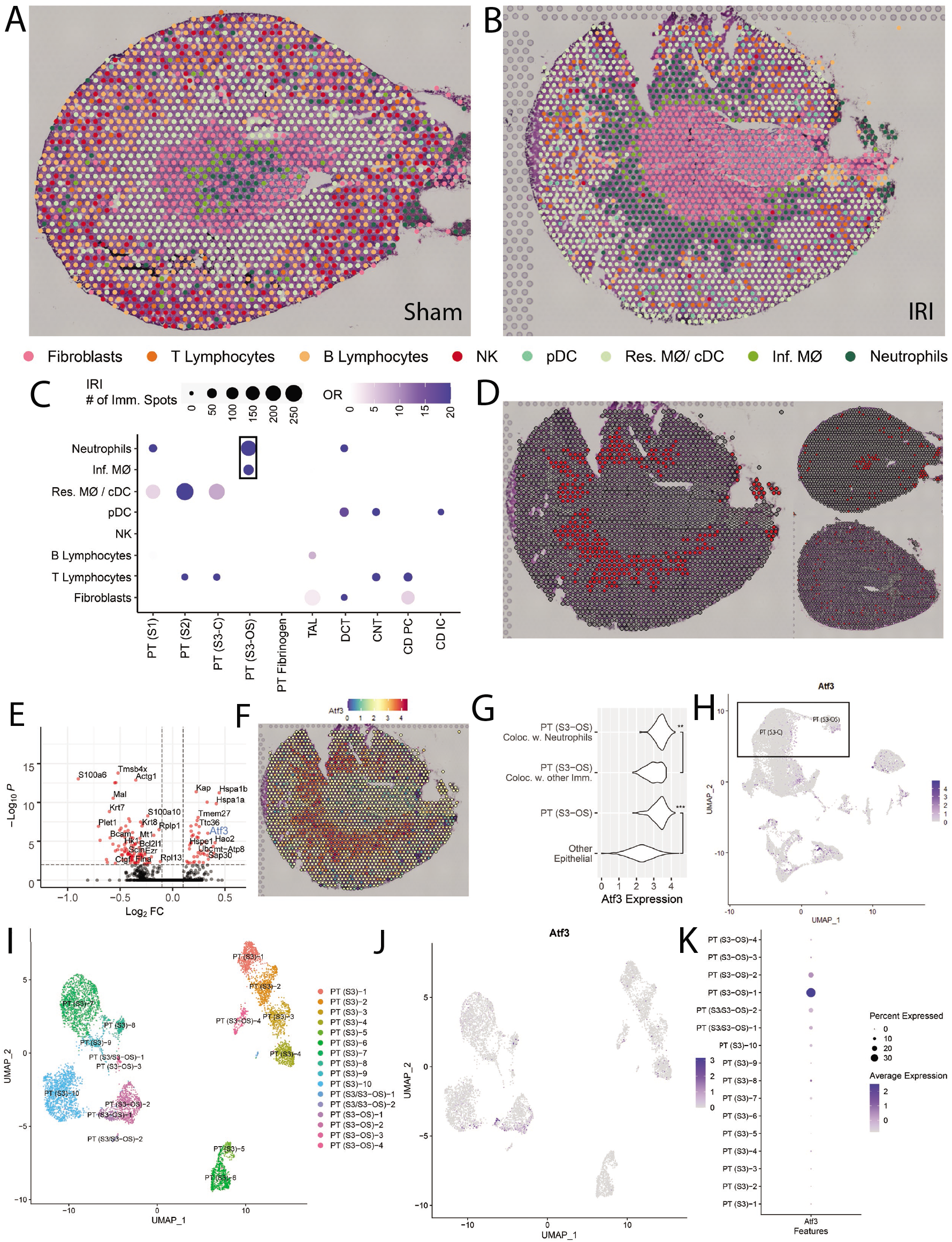
Colocalization of immune clusters in the Ischemia Reperfusion Injury model. (**A-B**) Transfer of selected single cell immune clusters overlaid upon the spatial transcriptomics sections for the Sham and Ischemia Reperfusion Injury (IRI) models, respectively. (**C**) Dot plot presenting the odd ratio of colocalization of a pair of clusters in IRI when compared to Sham. (**D**) Highlight of Neutrophils in IRI (left), Sham (top-right) and Cecal Ligation Puncture (CLP, bottomright). (**E**) Volcano plot presenting the Differentially Expressed Genes (DEGs) between the PT (S3-OS) spots colocalizing with Neutrophils (right) and the ones colocalizing with other immune clusters in IRI (left). (**F**) Feature plot presenting the expression levels of Atf3 in IRI. (**G**) Violin plot comparing the expression distribution of Atf3 in selected clusters (** - p<10^−9^, *** - p<10^−15^). (H) Feature plot of the single cell data presenting the expression of Atf3 with PT (S3-C) and PT (S3-OS) clusters highlighted. (I) UMAP of the PT (S3-C) and PT (S3-OS) subclusters in the single cell data. (J) Feature plot of Atf3 with its expression on the single cell subclusters. (K) Dot plot comparing the expression of Atf3 in all subclusters. (Abbreviations: PT – Proximal Tubule; S1, S2, S3 – Segments of proximal tubule; S3-C – Cortical section of S3; S3-OS – Outer Stripe section of S3; TAL – Thick Ascending limb; DCT – Distal Convoluted Tubule; CNT – Connecting Tubule; CD – Collecting Duct; PC – Principal Cells; IC – Intercalated Cells; NK – Natural Killer Cells; pDC - Plasmacytoid Dendritic Cells; cDC - Conventional Dendritic Cells; Res. MΦ – Resident Macrophages; Inf. MΦ – Infiltrating Macrophages). Each spot is 55 μm in diameter.

**Figure 7:**
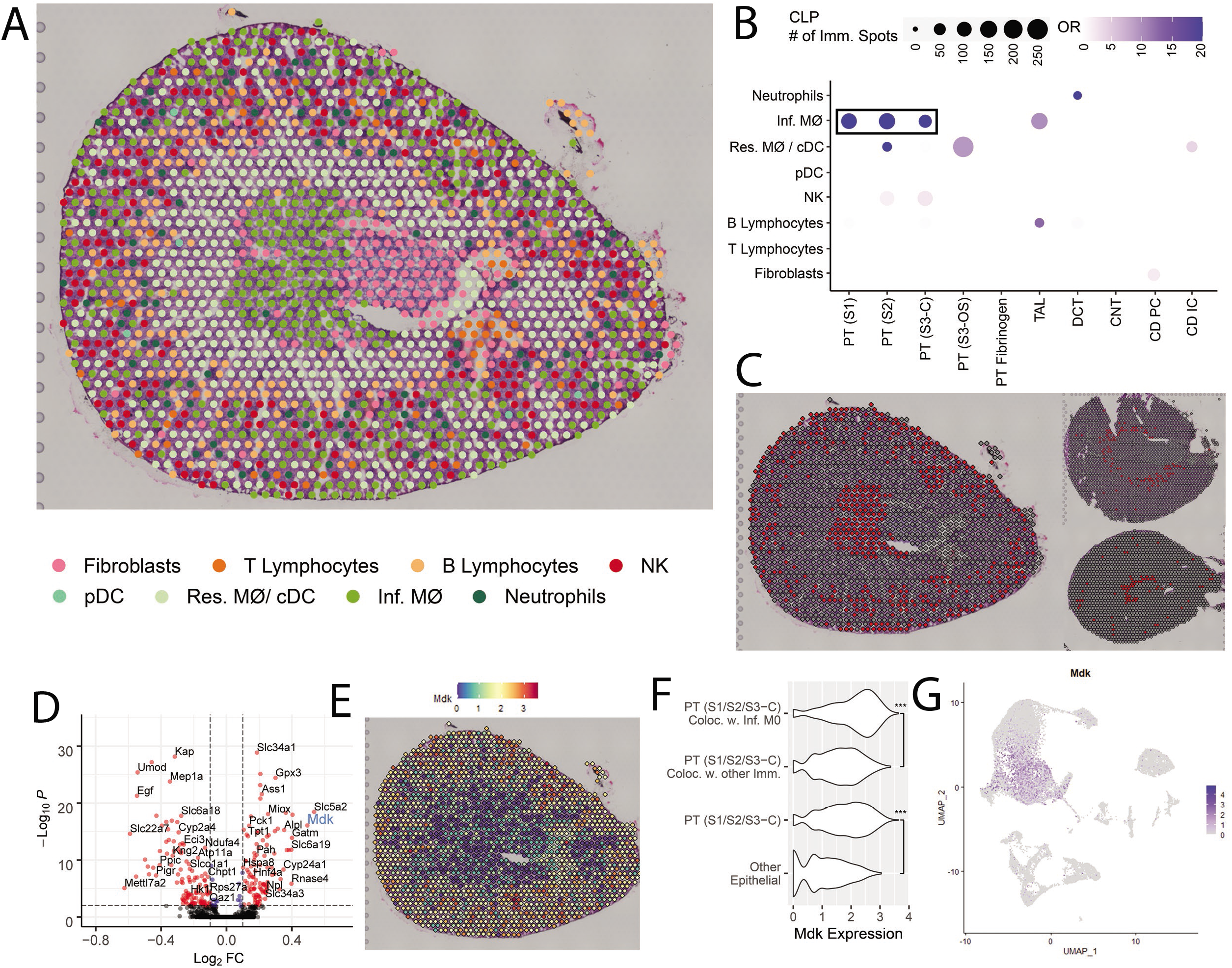
Colocalization of immune clusters in the Cecal Ligation Puncture model. (A) Selected single cell immune clusters transferred over Cecal Ligation Puncture (CLP) spatial transcriptomics. (B) Dot plot presenting the odds ratio of colocalization of a pair of clusters in CLP when compared to Sham. (C) Highlight of Infiltrating Macrophages in CLP (left), Sham (top-right) and Ischemia Reperfusion Injury (bottom-right). (D) Volcano plot presenting the Differentially Expressed Genes (DEGs) between the PT (Sl/S2/S3-C) spots colocalizing with Infiltrating Macrophages (right) and the ones colocalizing with other immune clusters in CLP. (E) Feature plot presenting the expression levels of MdK in CLP. (F) Violin plot comparing the expression distribution of Mdk in selected clusters (*** - p<10-15). (G) Feature plot presenting the expression of MDK in the single cell data. (Abbreviations: PT – Proximal Tubule; S1, S2, S3 – Segments of proximal tubule; S3-C – Cortical section of S3; S3-OS – Outer Stripe section of S3; TAL – Thick Ascending limb; DCT – Distal Convoluted Tubule; CNT – Connecting Tubule; CD – Collecting Duct; PC – Principal Cells;

### Colocalization of immune and epithelial cell transcriptomic signatures in spots of the ischemia reperfusion model

Figures 6A and 6B present the spatial distribution of immune cell and fibroblast clusters mapped onto the sham and IRI models, respectively. A transfer score was quantitated for each scRNAseq cluster signature for every ST spot. Only the immune cell with the highest ranked transfer score is displayed. Fibroblasts were included in the analysis as a control, i.e. the immune cell transfer score had to at least exceed that of a fibroblast to be mapped to the histologic section.

These new immune cell spot identities were then compared to the original epithelial cell type spot identities mapped in Figure 5.

To evaluate the colocalization of immune and epithelial cells, an odds ratio of colocalization was calculated across all ST spots (Figure 6C, Supplemental Table S8) as compared to the sham mouse. The strongest association was found between the proximal tubule S3 spots of the outer stripe (PT (S3-OS)) with the neutrophil and infiltrating macrophage signature. The remaining significant cluster pairs possessed lower odds ratios or were indicative of resident macrophage colocalization. We elected to explore the relationship between the PT (S3-OS) and neutrophil migration.

The neutrophil scRNAseq cluster mapped specifically to ST spots of the outer stripe of the medulla in the IRI model (Figure 6D). Such co-clustering was not apparent in the sham and CLP models. In the IRI model, we then queried DEGs between the PT (S3-OS) spots that colocalized with neutrophils and the PT (S3-OS) spots that colocalized with any other immune cell type (Figure 6E, Supplemental Table S9). Among the DEGs was Atf3, a regulator of neutrophil migration (24). The expression of Atf3 specifically colocalized to the outer stripe of the medulla (Figure 6F). The expression of Atf3 was both higher in the PT (S3-OS) that colocalized with neutrophils than those PT (S3-OS) that did not and the expression of Atf3 was higher in the PT (S3-OS) than all other epithelial cell types (Figure 6G).

Since a subpopulation of PT S3 tubules expressing Atf3 was detected, the expression of Atf3 was assessed in the single cell data (Figure 6H). The presence of small tightly grouped cells with high expression of Atf3 suggested the presence of possible meaningful subpopulations. We then re-clustered the cortical and outer-stripe clusters of PT S3 to better identify the subpopulation equivalent to the one detected in ST. Figure 6I shows the UMAP of the subclusters, while Figure 6J, the expression of Atf3 in those cells. Finally, Figure 6K shows we were able to identify a subpopulation of cells originating from outer stripe that highly expresses Atf3. Thus, the ST data facilitated the discovery of a new cell type within the scRNAseq dataset: a PT S3 cell which is signaling neutrophils.

As a control, we identified S100a6 as a significantly down-regulated gene in the PT (S3-OS) cluster that co-localized with neutrophils (Supplemental Figure S3). This gene was not highly expressed in the outer stripe by spatial transcriptomics. In the scRNAseq dataset, S100a6 was expressed across most immune cell types, including neutrophils.

### Colocalization of immune and epithelial cell transcriptomic signatures in spots of the cecal ligation puncture model

In a corresponding analysis, we then investigated the transfer of the scRNAseq immune cell clusters over the CLP section (Figure 7A). The odds ratio of colocalization (Figure 7B, Supplemental Table S8) indicates macrophage signature colocalization throughout the cortical segments of proximal tubules. Chil3+ and Ear2+ infiltrating macrophage clusters were merged to increase detection power. The spots containing a highest-ranking macrophage transfer score (i.e. strongest macrophage expression signature components) are highlighted in the three models (Figure 7C) and suggest that macrophage infiltration was most pronounced in the cortex of the CLP mouse as compared to the sham or IRI model. DEGs were assessed between PT spots in the CLP model that colocalized with infiltrating macrophage clusters versus those PT spots that colocalized with any other immune cell or fibroblasts (Figure 7D, Supplemental Table S9). Among the top DEGs associated with macrophage colocalization was Mdk, a gene encoding a growth factor associated with macrophage recruitment (25), which was more highly expressed in the outer cortex of the CLP section (Figure 7E). The expression of Mdk was higher in cortical PT spots that colocalized with infiltrating macrophages as compared to cortical PT spots that did not colocalize with macrophages and as compared to all other epithelial cell types (Figure 7F).

In contrast to Atf3, the expression of Mdk in the scRNAseq dataset (Figure 7G) did not indicate clear subpopulation of Mdk expressing PT cells, but was instead diffusely expressed across the proximal tubule. In addition to the colocalization of the macrophage signature with the proximal tubule signature, a smaller, but still significant association was observed between the NK cell signature and the proximal tubular signature in the CLP model.

### Visualization of immune cells by multiplexed immunofluorescence imaging of the murine injury models

To validate the localization of immune cells in each model, we performed CODEX multiplexed immunofluorescence imaging on sections from the same animal (26). Figure 8 reveals the immune cell distribution in the sham, IRI, and CLP models. Supplemental Figure S4 presents the gating strategy employed in CODEX. In comparison with the sham model, neutrophils (Ly6G+ and CD11b hi) were found to predominantly infiltrate the outer stripe of the IRI section (Figure 8D). The percentage of neutrophils and infiltrating macrophages in each approximate region of the kidney (papilla, inner medulla, outer stripe of the medulla, and cortex) was quantitated (Figure 8D-F). In the IRI model, 52.2% of all infiltrating neutrophils localized to the outer stripe. This finding is consistent with the localization of neutrophil cluster ST spots to the outer stripe (Supplemental Table S10).

**Figure 8:**
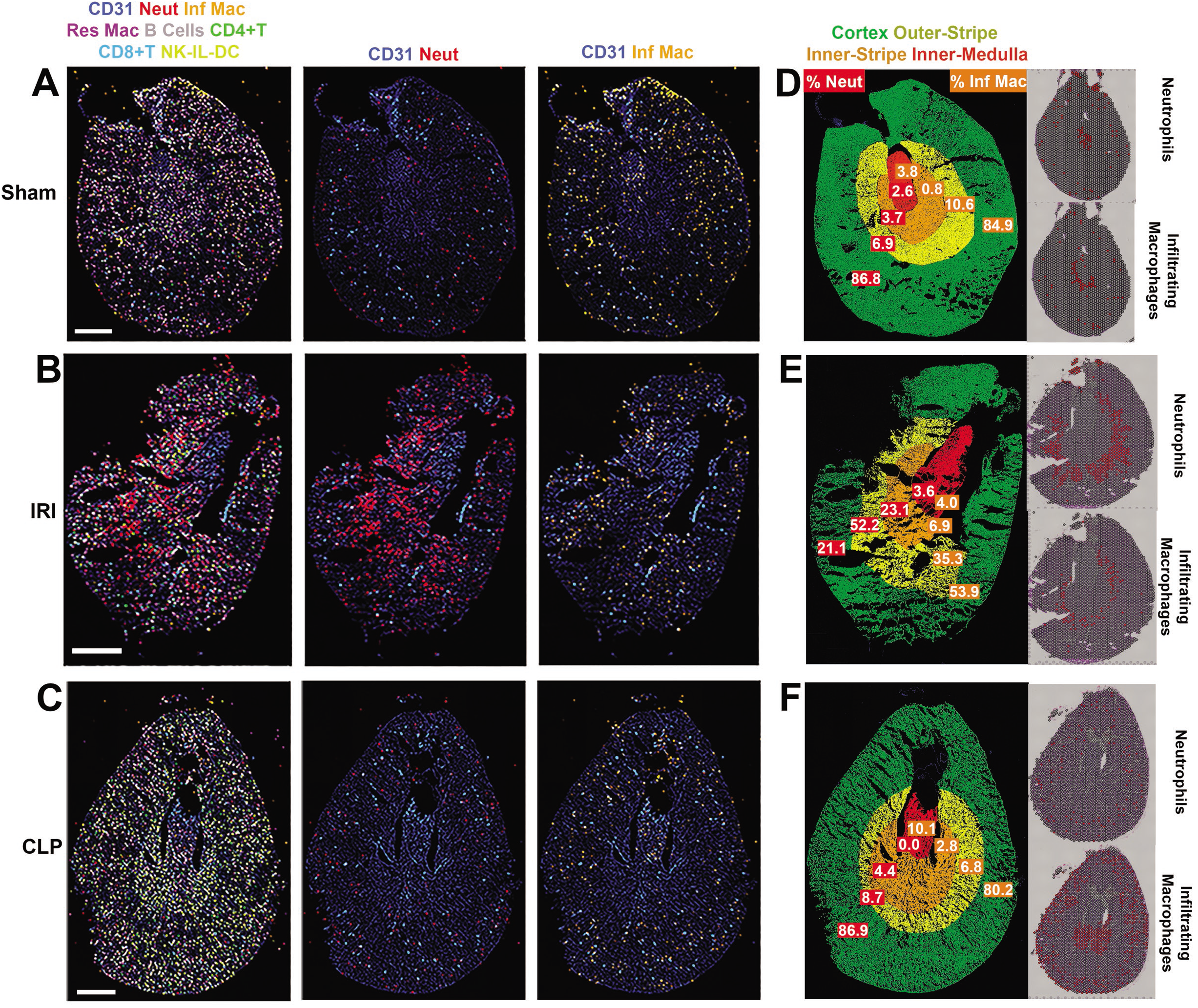
Multiplexed imaging of proteins in toto with CODEX validates the localization of immune cell clusters inferred by Spatial Transcriptomics. CODEX imaging for kidney sections from Sham, Ischemia Reperfusion Injury (IRI) and Cecal Ligation Puncture (CLP) are shown in (A), (B) and (C), respectively. In the left column for all sections, spatial mapping of various immune cells (neutrophils, infiltrating and resident macrophages, B cells, CD4+ and CD8+ T cells and NK/IL/DC cells) are displayed using colored overlays, and CD31 staining is included for context. The definition of each cell type based on the presence and absence of markers is detailed in Supplemental Figure 4. The second column shows only CD31 and neutrophils (Neut, red), and the third column displays infiltrating macrophages (Inf Mac, orange). D-F show the distribution of neutrophils and infiltrating macrophages in specific regions of the kidney for Sham, IRI and CLP, respectively. The cortex, outer stripe of the medulla, inner medulla, and papilla regions were identified based on structural landmarks and annotated using region of interest (ROI) tool in ImageJ. The corresponding spatial transcriptomic signature for neutrophils and infiltrating macrophages is shown on the right side for each specimen.

In the CLP model, 80.2% of infiltrating macrophages (Ly6G- and CD11b hi) localized to the outer cortex (Figure 8F). The infiltrating macrophage signature dominated and colocalized with proximal tubule epithelial spots in the spatial transcriptomic analysis (Figure 7C), and the distribution of these macrophages localized to the outer cortex of the kidney by immunofluorescence. However, no increase in macrophage quantity was observed in the CLP model as compared to the sham model. We speculate that the activated macrophages in the CLP model contributed to the transcriptomic signature to a greater extent than the macrophages in the sham model.

A second finding in the CLP model was the infiltration of natural killer cells in the cortex and outer stripe of the kidney. Cells consistent with Natural Killer cells (CD45+, CD11b-, B220-, CD3-, CD4-, CD8-) localized to these regions in the CODEX assay. Further, the NK cell infiltration aligned with ST colocalization of NK cells with PT S2 and PT S3 expression signatures in the CLP model (Figure 7B). In summary, the ST signatures uncovered in the IRI and CLP models were supported by the immunofluorescence data.

## Discussion

In this work, we localize the transcriptomic signature of various immune cells to spatial transcriptomic spots of known renal epithelial cells in two models of AKI. Signatures from immune and epithelial cells were colocalized to identify possible chemotactic molecules of neutrophils and infiltrating macrophages in the IRI and CLP models respectively. Further, the spatial transcriptomic analysis enhanced our understanding of the single cell data, by detecting a subpopulation of injured proximal tubule cells with Atf3 expression which may be responsible for neutrophil chemotaxis. To accomplish this, we first optimized a workflow for the interrogation of human and murine kidney tissue with spatial transcriptomics. The technology facilitated the identification of key renal cell types and regions, including most epithelial, endothelial, and stromal cell types. The specificity of cell type mapping was improved by concomitant snRNAseq or scRNAseq cluster analysis. The technique appears robust, revealing strong concordance (>97%) between the transcriptomic signature of a given spot and its underlying histopathology. Supporting data is provided that acute kidney injury (AKI) does not affect all regions of the kidney uniformly. The early effects of acute tubular necrosis in the IRI model led to regional changes in the transcriptomic signature at the 6 h timepoint.

Single cell sequencing has rapidly enhanced our knowledge of the kidney’s expression signature and its cell types (27), thereby helping to uncover the pathophysiology of a variety of conditions including early diabetic nephropathy (28), the composition of a Wilm’s tumor (29), and allograft rejection (5). Similar studies in the mouse have facilitated query of murine disease state models (30) and the response to water deprivation (31). In addition, scRNAseq and snRNAseq are foundational technologies for the generation of a kidney cell atlas and detection of novel cell types in the human (2) and the mouse (4, 32, 33). scRNAseq has also aided in our understanding of kidney organ development, as studies performed in organoids have shown expression of disease markers in early glomerular cells (34) and revealed that organoid expression patterns maintain strong agreement with fetal kidney expression, including lineage differences and growth signatures (35, 36). Single cell analysis has also aided in understanding the development of fetal kidney (37–40) and myofibroblast origin (41).

In our study, we used the snRNAseq and scRNAseq datasets to effectively expand the breadth of cell types mapped to kidney tissue by ST. For example, the snRNAseq clustering in human kidney allowed us to differentiate interstitium spots into dominant signatures of endothelial cells, fibroblasts and VSMCs. In the mouse, we were able to detect multiple clusters not present in the unsupervised ST dataset, particularly immune cells. Because there are often up to 10,000 single nuclei or cells in a given sample, the scRNAseq and snRNAseq technologies have improved sample size to detect less represented cell types. Furthermore, the scRNAseq data is cell-specific rather than representative of a 55 μm spot with multiple underlying cells.

Multiple ST platforms exist (42), including both slide seq and the 10x Visium platform (6). ST has been used to establish atlases in other organs such as mouse brain (6) and the human heart (43). In the context of disease, the spatial heterogeneity of prostate cancer (44) and melanoma in lymph nodes (45) has been described, along with the progression of amyotrophic lateral sclerosis in mice (46). The advantages of the 10x Visium platform, used here, include direct visualization of the underlying histology along with a higher sensitivity for gene detection. Other *in situ* mRNA capturing technologies maintain a higher resolution, but lower sensitivity for gene expression detection. These alternatives often require the histology image from a sequential slice (47, 48) or capture smaller tissue areas (49). Other approaches to determine the spatial distribution of mRNA, such as *in situ* sequencing (50) or sequential *in situ* hybridization (51) are expensive, but important ways to visualize the transcriptomic signature with spatial resolution.

The main limitation of the 10x Visium Spatial Gene Expression platform is its resolution with a spot size of 55 μm, which will overlay multiple cells. For instance, Slide seq has improved cellular resolution, but has not been optimized for direct visualization on the same histopathology H+E stained specimen (42). In this work, we chose to examine the 10x Visium platform as it had the ability to create an atlas of the kidney with direct mapping of expression onto the renal structures a pathologist would assess, thus being more readily translatable. We overcame the limitation in resolution by deconvoluting each spot with transfer scores, ranking the contribution of each scRNAseq cluster cell type to each transcriptomic spot signature. The limitations of this technology are further counterbalanced by its deep signature (approximately 3,000 genes detected and 10,000 reads per spot with over 16,000 genes per cluster) and the ability to directly visualize the signature over a histologic image. Thus, this technique affords more than simply identifying the proximity of localized scRNAseq cells, it can actually identify distinct molecular signatures (as in the case of IRI) and correlate these with histology. We leveraged the advantages of the 10x platform to assess regional distribution of kidney injury expression changes in both murine models. We counted total nuclei in size-matched regions which correlated with total expression in the IRI model in those regions. An additional limitation is that each capture area has about 5,000 spots; therefore, even if the sample covers the capture area completely, the number of spots may not be enough to cluster and differentiate all cell types. As stated above, we were able to effectively use scRNAseq to expand the breadth of cell types mapped.

In this work, we also present a technique to deconvolute and colocalize transcriptomic signatures of multiple cell types within a single spot. This approach was employed to detect immune cell localization and uncover potential chemotactic signals. This form of deconvolution has been used to classify secondary contributors to a spot’s expression signature, in a similar way to what has been done in the heart (52). The immune cell population of the kidney has been extensively studied using scRNAseq in healthy subjects and in the context of diabetic nephropathy and lupus nephritis (3, 53, 54). In our study, we unite scRNAseq with ST to study the immune cell distribution in known murine models of AKI. Much is known of the role of immune cells in these common models of AKI. It has been shown in IRI models, that infiltration of neutrophils occurs in the outer stripe of the medulla (55, 56). Further, macrophage colocalization in the S1 segment of proximal tubules has been found protective in sepsis models (57, 58). Thus, the localization of these immune cells in the AKI models is not the novelty, instead, it is the capture of gene signatures underlying this colocalization. We used 10x Visium to predict the spatial distribution of immune cell types in the kidney and identify potential chemotactic factors associated with particular epithelial cells. The regional distribution of these cells was validated with CODEX multiplexed immunofluorescence. By design, this work is descriptive. Future investigations are needed to understand the cause-and-effect relationship of these immune cell signals in the kidney.

### Conclusions and future directions

In summary, we present the spatially mapped transcriptomic signature of AKI in murine models and show how this methodology can be applied to human kidney tissue. In the future, this technology may assist renal pathologists in their interpretation of kidney biopsy specimens. The ability to link upregulation of a particular injury gene to a particular nephron structure is vitally important. This might allow improved classification of human AKI. Future endeavors will seek to quantitatively define the contribution of each cell type signature to each spot and to apply this methodology to diseased human kidney tissue.

## Methods

### Human tissue, data source, and snRNA sequencing

Publicly available snDrop-seq RNA sequencing data were acquired for six Kidney Precision Medicine Project (www.KPMP.org) samples and nine additional samples from the Washington University Kidney Translational Research Center (GEO, GSE121862). Sequencing data of nuclei for the samples were subjected to quality control metrics as previously described (2). Using Seurat 3.1 and PAGODA2 (https://github.com/hms-dbmi/pagoda2), nuclei were reclustered and displayed as a uniform manifold approximation and projection (UMAP). A single human reference nephrectomy (female aged 59 without histologic evidence of kidney disease) was acquired from the Biopsy Biobank Cohort of Indiana (BBCI) Eadon, AJKD). This study was approved by the Institutional Review Board (IRB # 1906572234) of Indiana University.

### Murine models

Animal experiments and protocols were approved by the Indiana University Animal Care and Use Committee. From age-matched 8-10 week old 129/ SvEv mice (Taconic Biosciences, Albany, New York), tissues were acquired from a sham mouse in which the abdomen was opened and sutured back and mice which underwent ischemia-reperfusion injury (IRI) or cecal ligation and puncture (CLP). In the IRI model (59), both renal pedicles were exposed and clamped for 22 minutes through a midline incision then released. In the CLP model (58), the cecum was ligated and punctured with a 25 gauge needle. Kidneys were excised upon sacrifice 6 hours after each procedure and frozen in Optimal Cutting Temperature (OCT) compound. The presence or absence of acute kidney injury (AKI) was assessed on the H+E histology image and in a consecutive periodic acid-Schiff stained section. As expected, blood urea nitrogen and creatinine measurements are not elevated in either the IRI and CLP models at the 6 h timepoint.

### Slide preparation and imaging

Slide preparation (CG000240_Demonstrated_Protocol_VisiumSpatialProtocols_TissuePreparationGuide_Rev_A, 10X Genomics) and imaging were conducted according to Visium Spatial Gene Expression protocols (CG000241_VisiumImagingGuidelinesTN_Rev_A, 10X Genomics). Frozen transverse 10 μm sections from the human nephrectomy or each murine model were placed within the etched frames of the capture areas on the active surface of the Visium Spatial Slide. Tissue sections were fixed in methanol and stained with hematoxylin and eosin (H&E). Brightfield images of stained sections in the fiducial frames were collected as mosaics of 10x fields using a Keyence BZ-X810 microscope equipped with a Nikon 10X CFI Plan Fluor objective.

### mRNA extraction and sequencing

mRNA extraction, library preparation, and sequencing were conducted according to the Visium spatial protocols. Stained tissue sections were permeabilized for 12 minutes, mRNA was released to bind oligonucleotides on the capture areas, followed by reverse transcription, second strand synthesis, denaturation, cDNA amplification, SPRIselect cDNA cleanup (CG000239_VisiumSpatialGeneExpression_UserGuide_RevD, 10X Genomics), and then the cDNA libraries were prepared and sequenced on an Illumina NovaSeq 6000 with 28bp+120bp paired-end sequencing mode.

### Murine single cell isolation, library preparation, and sequencing

Sham and IRI murine kidneys were transported on ice, minced, and dissociated using the Multi-Tissue Dissociation Kit 2 and a dissociator tube rotator (GentleMACS, Miltenyi Biotec). Samples were prepared according to protocol with modifications: 10 mL of RPMI1640 (Corning) and 10% BSA (Sigma-Aldrich) was added to the mixture, then filtered, centrifuged at 300 g for 5 minutes, and the cell pellet was re-suspended in 1 mL of lysis buffer (Sigma). Annexin V dead cell removal was performed using magnetic bead separation after washing. The pellet was resuspended in RPMI1640 and BSA 0.04%. A final concentration of 1 million cells/mL with over 80% viability was achieved. In a single cell master mix with lysis buffer and reverse transcription reagents, the sample was processed according to the Chromium Single Cell 3’ Reagent Kits V3 protocol (CG000183_ChromiumSingleCell3_v3_UG_Rev-A, 10X Genomics). cDNA was synthesized and libraries were prepared. Sequencing was performed on the Illumina NovaSeq6000 with 28bp+91bp paired-end mode.

scRNA seq was not performed for CLP mice. Instead, publicly available kidney scRNAseq data was downloaded from GEO (GSE154107) for mice exposed to lipopolysaccharide (LPS, 5 mg/kg) or vehicle and sacrificed at 4 hours as previously described (21).

### Murine single cell data processing

Data from four scRNAseq experiments (2 sham, 1 IRI, 1 LPS) were processed with Cell Ranger 2.1.0, and Seurat 3.1 in R (60). Raw base call files were demultiplexed from FASTQ files and aligned to the mm10 murine genome using STAR (Dobin 2013). Cells were removed based on a mitochondrial content higher than 50%, less than 200 unique genes, and the lower 10% and top 5% percentiles of unique genes distribution on each mouse. Each mouse dataset was independently scaled and normalized and then merged to create a single cell reference. After defining the clusters with a resolution of 0.65, the immune, interstitium, and endothelial clusters were reclustered with a higher resolution of 1.4.

### Spatial transcriptomics expression analysis

Mapping and counting were performed using Space Ranger 1.0.0 with the reference genome GRCh38 3.0.0 or mmu10 provided by 10x genomics. Space Ranger aligns the barcodes in each read with a 55 μm “spot” coordinate relative to the fiducial frame, associating read counts with the image. After generating counts, Space Ranger clusters spots using a graph-based clustering algorithm where a nearest neighbor network is built in a Principal Components space and a Louvain Modularity Optimization algorithm selects the modules of highly connected spots. Finally, Space Ranger calculates differential expression between the clusters to identify differentially expressed genes (DEGs) using the sSeq method with a t-test for spots with low counts and edgeR method with asymptotic beta test for cells with high counts. The feature plots present expression levels are normalized by Space Ranger. After mapping of the three mouse samples by Space Ranger, the data was processed in Seurat 3.1. The data was normalized by SCTransform and merged to build a unified UMAP. Glomerular spots in the Sham model were manually annotated based on the composite reference UMAP. Dot plots show only above average expression.

Seurat was used to transfer cluster labels from snRNAseq and scRNAseq to ST spots in humans and mice, respectively. The integrated murine scRNAseq clusters were transferred to each murine ST sample. The transfer procedure generates a probability score for each spot and its association with a given scRNAseq or snRNAseq cluster. The spot is assigned to the cluster with the highest score and mapped back to the ST sample image. For deconvolution analyses, dominant non-immune clusters were suppressed in the analyses and the immune cell types were with the highest probability transfer score were mapped on each mouse.

### Correlation with histology

In the human nephrectomy, five fields were selected at random, each composed of 237 to 245 spots, representing 40.0% of all spots mapped to the nephrectomy tissue. The histology underlying each spot was assessed and the percentage of concordance was determined. Spots were only counted if completely in frame. A spot was considered in alignment if any portion overlapped histology consistent with cluster identity.

### Multiplexed CODEX immunofluorescence

Murine tissue sections of 10 μm were acquired from the same OCT block as the spatial transcriptomics sections and were collected on coverslips coated in poly-L-lysine. Sections were prepared following the protocol provided by the manufacturer Akoya Biosciences, which is also described in detail by Goltsev et al (26). Tissue retrieval was conducted with a 3-step hydration process. After hydration, tissues were fixed for staining. An antibody cocktail was made using the antibodies listed below and dispensed among the three tissues. After staining, another fixation step was performed to ensure adherence of the tissue to the slide, in preparation for washing cycles during imaging. Sections were imaged at 20x resolution using the fully automated CODEX system (Akoya Biosciences) and a Keyence BZ-X810 slide scanning microscope. Samples were processed, analyzed, and visualized in the CODEX-MAV software, a plugin for FIJI/ImageJ. The following antibodies and their barcodes were used for staining (all from Akoya) : CD3-BX021 (17A2) – Cy5-RX021, CD4-BX026 (RM4-5) Atto 550-RX026, CD8a-BX029 (53-6.7) – Atto 550-RX029, CD11b-BX025 (M1/70) – Alexa FluorTM 488-RX025, CD31-BX002 (MEC13.3) – Atto 550-RX002, CD45-BX007 (30-F11)-Alexa FluorTM 488-RX007, CD45R/B220-BX010 (Ra3-6B2) – Alexa FluorTM 488-RX010, CD169-BX015 (3D6.112) – Cy5-RX015, Ki67-BX047 (B56) – Atto 550-RX047, Ly6G-BX024 (1A8)-Cy5-RX024, MHC II-BX014 (M5/114.15.2)-Atto 550-RX014. Signal specificity was validated by ensuring the disappearance of the signal in between the imaging cycles. Gating strategy was performed similar to commonly used approaches for profiling immune cells using flow cytometry (59). Total number of cells were reported, as well as percentages of total CD45+ leukocytes.

### Statistical analyses

The differentially expressed genes in each comparison were found with the Seurat function FindMarkers (Wilcoxon Rank Sum test). Pathway enrichment for those genes was performed with the R packages ReactomePA (61) and ClusterProfiler (62). An odds ratio was calculated to determine the likelihood of sets of scRNA-seq clusters colocalizing in the same ST spot. Fisher’s Exact tests were performed to test whether the odds ratio was greater than 1, and to verify the significance of spot distributions on regions of low expression.

## Author contributions

Conception, analysis, interpretation of data: RMF, MTE, TME, PCD, KSC, ARS, SW, LC, DJ, MJF, DB, TH. Conduct of experiments: YHC, MTE, ARS, SW, CG, XX, DJ, KWD. Drafting / revising the article: MTE, RMF, ARS, TME, PCD, CZ. Providing intellectual content: KJK, TH, TAS, KWD. Final approval of the version to be published: All authors.

## ACKNOWLEDGEMENTS

**General**: The authors would like to thank Sanjay Jain and Blue Lake, as well as the investigators of the Kidney Precision Medicine Project (www.kpmp.org) for their gracious support and advice. The 10X 3’ scRNA sequencing and 10X Visium Spatial 3’ transcriptome sequencing were performed in the Center for Medical Genomics at Indiana University School of Medicine. We would like to thank The Indiana Center for Biological Microscopy for imaging assistance.

## Funding

Support for this work was provided by the NIH/NIDDK K08DK107864 (M.T.E.); R01DK099345 (T.A.S.).

## Data and materials availability

Spatial transcriptomic data is archived in the Gene Expression Omnibus (GEO # pending). Single cell sequencing data is archived in (GEO # pending) and (GSE154107).

## DISCLOSURES

The authors have nothing to disclose.

---

**Supplemental Figure 1:**
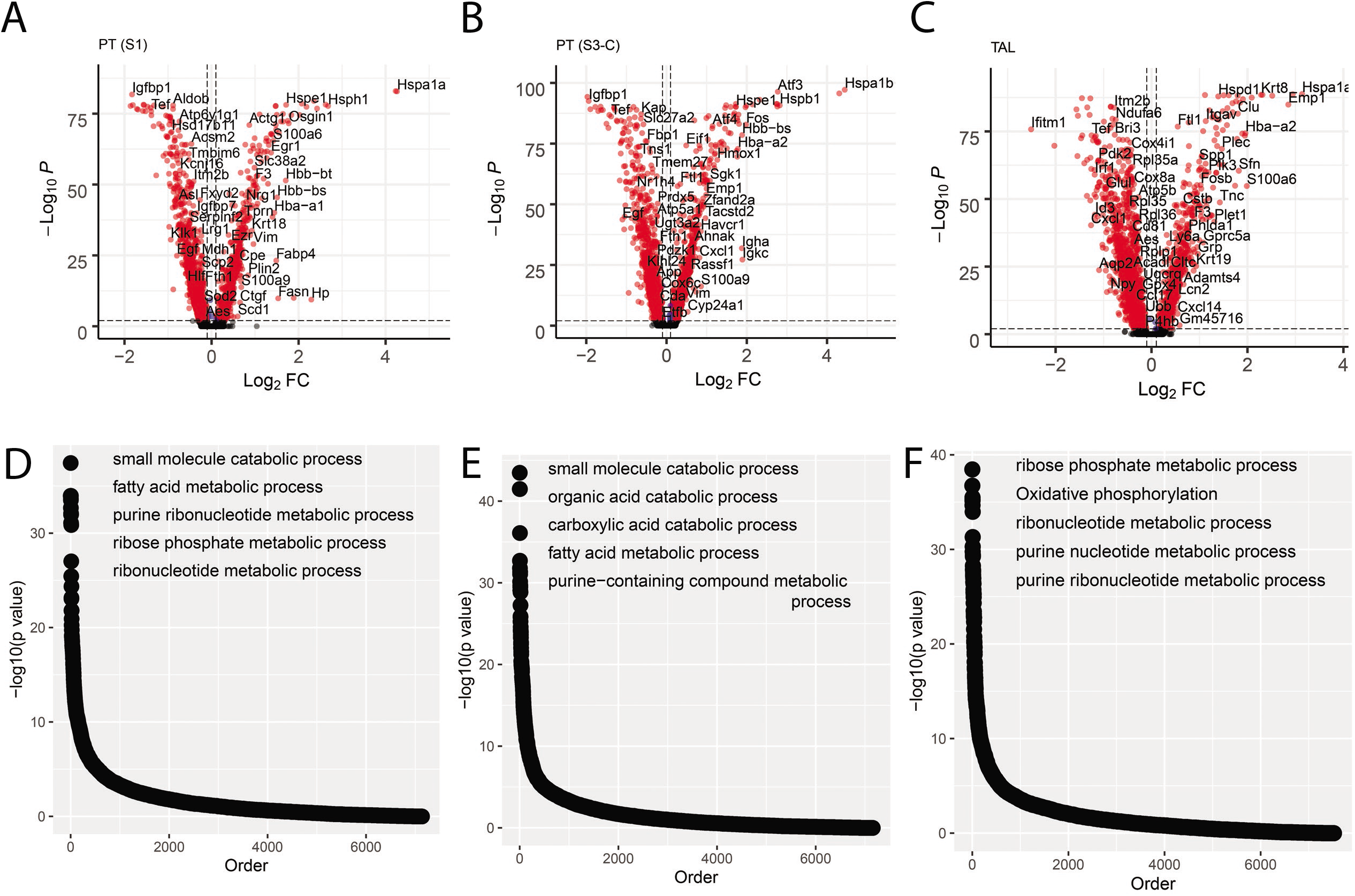
Differential expression between IRI and CLP models. Using clusters mapped from the scRNAseq dataset, (A-C) Differential expression in the single cell data between Ischemia Reperfusion Injury and Lipopolysaccharide murine kidneys is provided in selected clusters: PT (S1), PT (S3-C) and TAL, respectively. (D-F) Pathways enriched for the differentially expressed genes in the respective segment PT, (S1), PT (S3-C) and TAL, respectively. The top five pathways are annotated in the figure. (G) Odds ratio of overlap of differentially expressed genes between single cell and Spatial Transcriptomics. All Odds Ratio above zero are statistically significant. (Abbreviations: PT – Proximal Tubule; S1, S2, S3 – Segments of proximal tubule; S3-C – Cortical section of S3; S3-OS – Outer Stripe section of S3; TAL – Thick Ascending limb; DCT – Distal Convoluted Tubule; CNT – Connecting Tubule; CD – Collecting Duct; PC – Principal Cells; IC – Intercalated Cells; VSM – Vascular Smooth Muscle; NK – Natural Killer Cells; pDC – Plasmacytoid Dendritic Cells; eDC – Conventional Dendritic Cells; Res. MΦ – Resident Macrophages; Inf. MΦ – Infiltrating Macrophages).

**Supplemental Figure 2:**
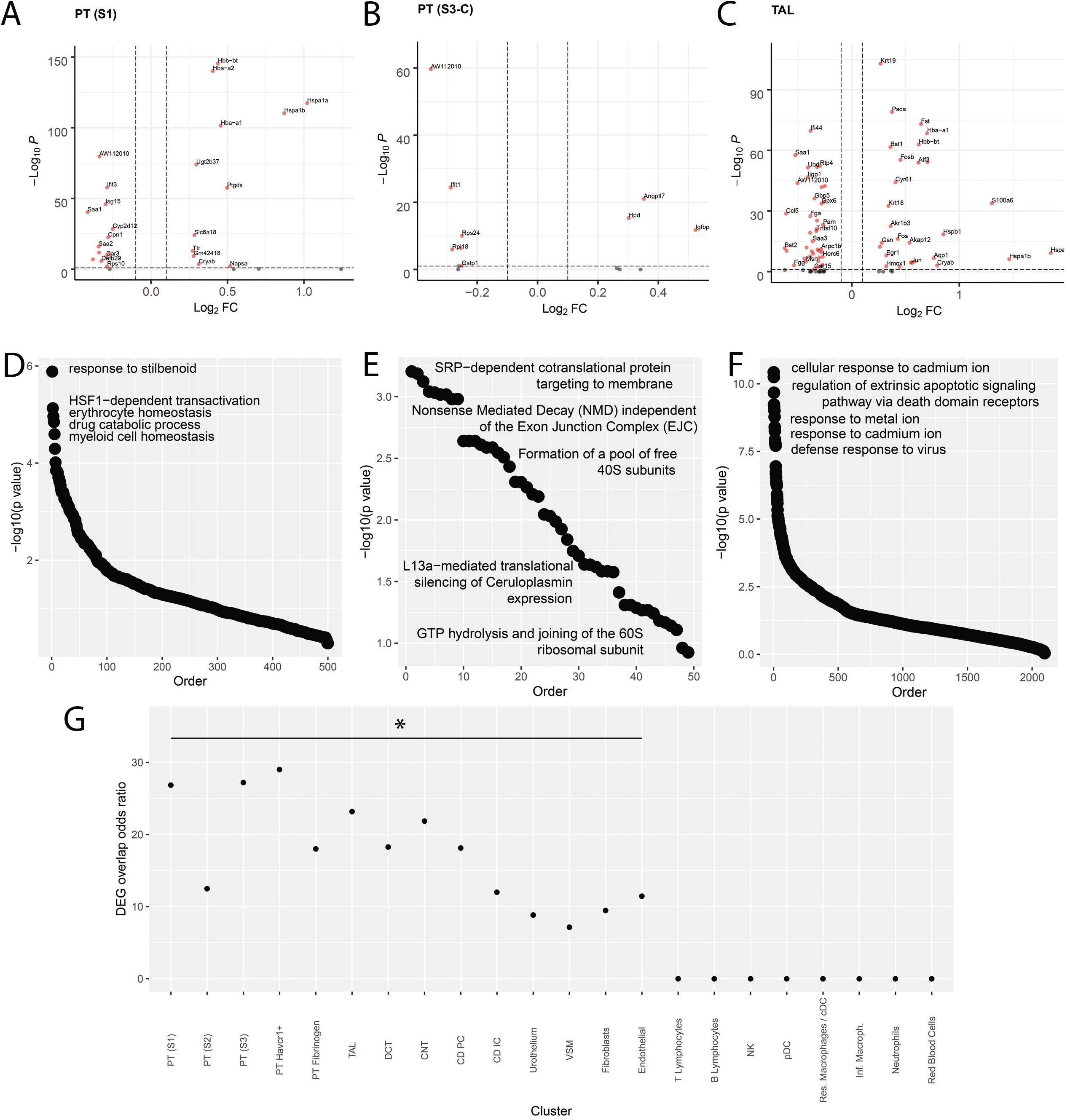
scRNAseq differential expression between IRI and LPS models. Using clusters mapped from the scRNAseq dataset, (A-C) differential expression in the single cell data between Ischemia Reperfusion Injury (IRI) and Lipopolysaccharide (LPS) murine kidneys by cluster, PT (S1), PT (S3-C) and TAL, respectively. (D-F) Pathways enriched for the differentially expressed genes in the respective segment PT, (S1), PT (S3-C) and TAL, respectively. The top five pathways are annotated in the figure. (G) Odds ratio of overlap of differentially expressed genes between single cell and Spatial Transcriptomics. All Odds Ratio above zero are statistically significant. (Abbreviations: PT – Proximal Tubule; S1, S2, S3 – Segments of proximal tubule; S3-C – Cortical section of S3; S3-OS – Outer Stripe section of S3; TAL – Thick Ascending limb; DCT – Distal Convoluted Tubule; CNT – Connecting Tubule; CD – Collecting Duct; PC – Principal Cells; IC – Intercalated Cells; VSM – Vascular Smooth Muscle; NK – Natural Killer Cells; pDC – Plasmacytoid Dendritic Cells; cDC – Conventional Dendritic Cells; Res. MΦ – Resident Macrophages; Inf. MΦ – Infiltrating Macrophages).

**Supplemental Figure 3:**
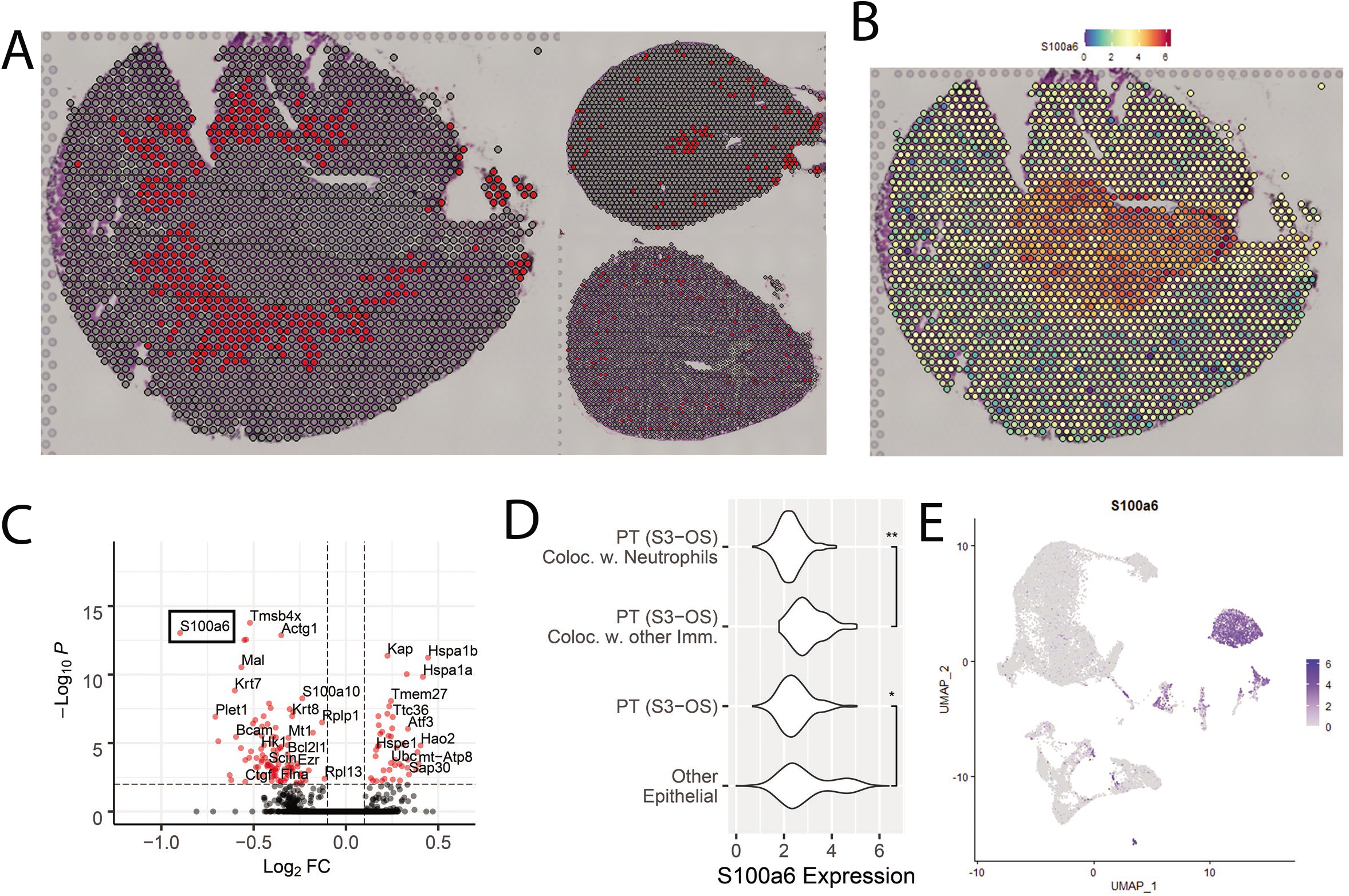
Analysis of S100a6 in colocalization of Neutrophils in Ischemia Reperfusion Injury. (A) Highlight of Neutrophils in Ischemia Reperfusion Injury (left), Sham (top-right) and Cecal Ligation Puncture (bottom-right) murine models. (B) Feature plot presenting the expression levels of Atf3 in Ischemia Reperfusion Injury. (C) Violin plot comparing the expression distribution of Atf3 in selected clusters (* - p<10-2, ** - p<10-9). (D) Feature plot presenting the expression of Atf3 in the single cell data. (E) Feature plot of S100a6 in the mouse single cell dataset. (Abbrevietion: PT (S3-OS) - S3 Segment of Proximal tubule located in Outer Stripe).

**Supplemental Figure 4:**
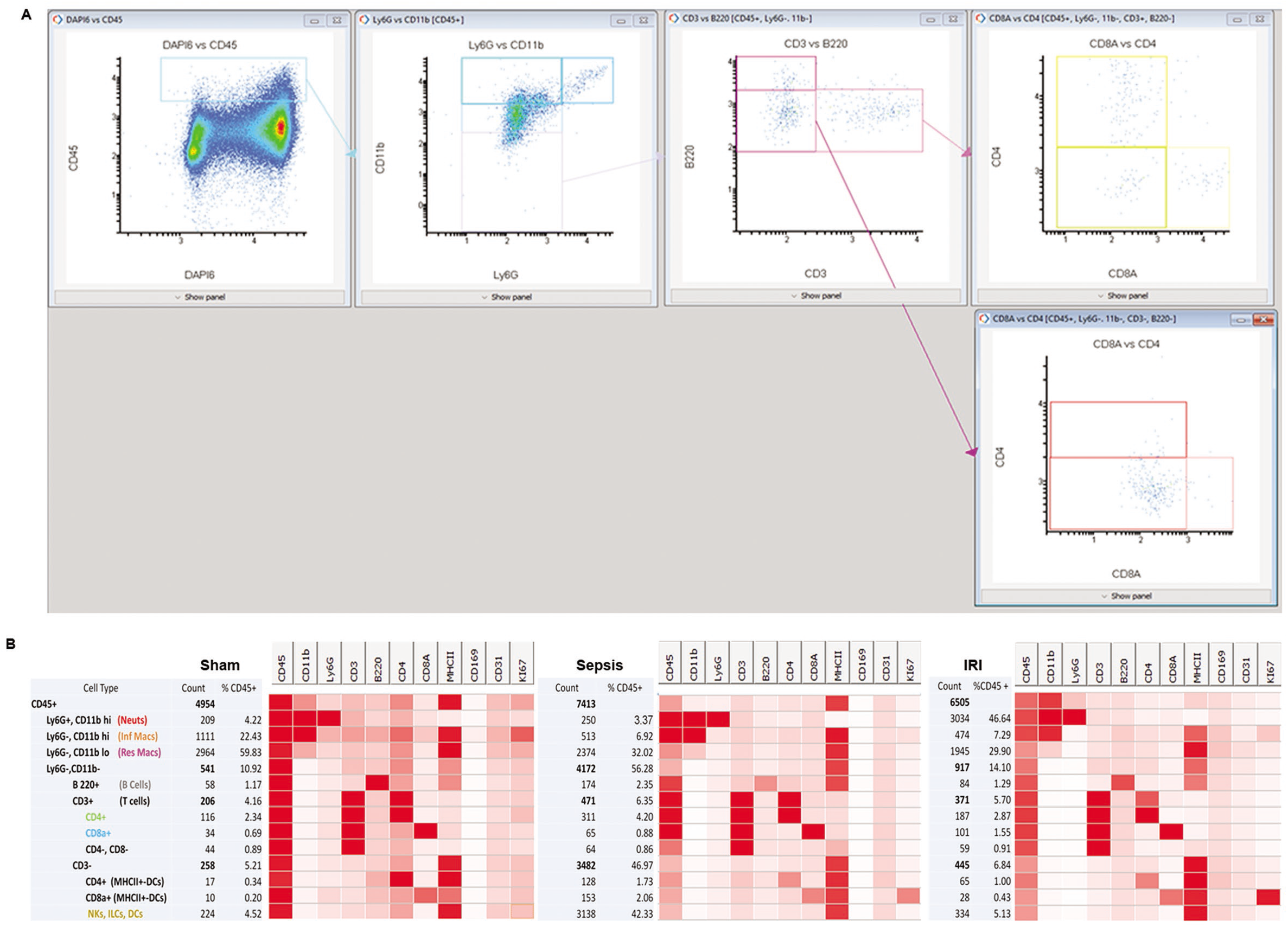
Gating strategy for CODEX analysis and multiplex profile of the immune cells. (A) Representative example of the gating strategy that was utilized for CODEX analysis. This strategy is based on flow cytometry profiling of immune cells. (B) Heatmaps depicting the relative distribution of median intensity fluorescence for each of the immune cell population in the three samples. The markers used are displayed in each column on the top, and the supervised analysis of immune cell populations based on the analysis used in (A) is shown in the left column, in a hierarchal pattern that follows the gating strategy. The number of cells in each population, as well as the percent of the total CD45+ cells are also shown. Of note, MHCII was not used in the supervised analysis and its level of expression in each cell type served for validation of the analytic strategy.

## Notes

### Competing Interest Statement

The authors have declared no competing interest.

